## Supplementary Information for "Super-resolution imaging in whole cells and tissues via DNA-PAINT on a spinning disk confocal with optical photon reassignment"

#### Supplementary Figures

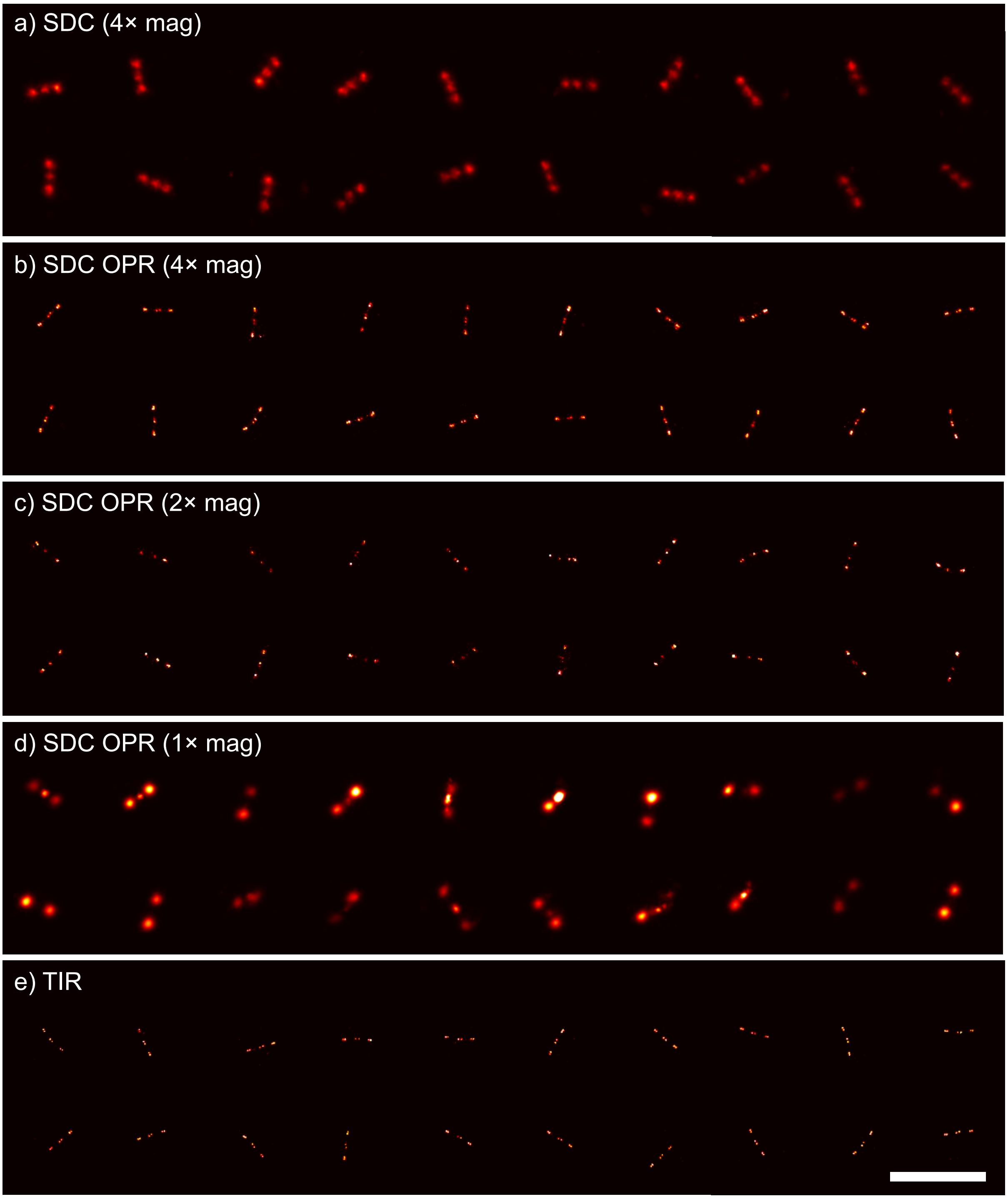


**Supplementary Figure 1**: Representative DNA origami from across the field of view of the measurement for different excitation illumination. (a) DNA origami structures imaged with spinning disk confocal (SDC) and 4× magnification. $\sigma_{SMLM}$: 7.0 nm. NeNa value: 11.5 nm.  Photon count per 300 ms: 869. (b) DNA origami structures imaged using 4× magnification with the spinning disk confocal microscope with optical photon reassignment (SDC-OPR), with deconvolution as described in methods. $\sigma_{SMLM}$: 1.4 nm. NeNa value: 2.3 nm. Photon count per 300 ms: 6990. (c) DNA origami structures imaged using 2.8× magnification with SDC-OPR, with deconvolution as described in methods. $\sigma_{SMLM}$: 1. nm. NeNa value: 3.1 nm. Photon count per 300 ms: 5614. (d) DNA origami structures imaged using 1× magnification with SDC-OPR, with deconvolution as described in methods. $\sigma_{SMLM}$: 8.3 nm. NeNa value: 16.7 nm. Photon count per 300 ms: 685. (e) DNA origami structures imaged using total internal reflection (TIR) illumination. $\sigma_{SMLM}$: 1.0 nm. NeNa value: 2.7 nm. Photon count per 100 ms: 13779. DNA origami sequence is shown in the methods, along with sample preparation, and imaging conditions are described in Supplementary Table 1. Scale bar is 500 nm. All DNA origami sample imaging were repeated in two independent experiments.

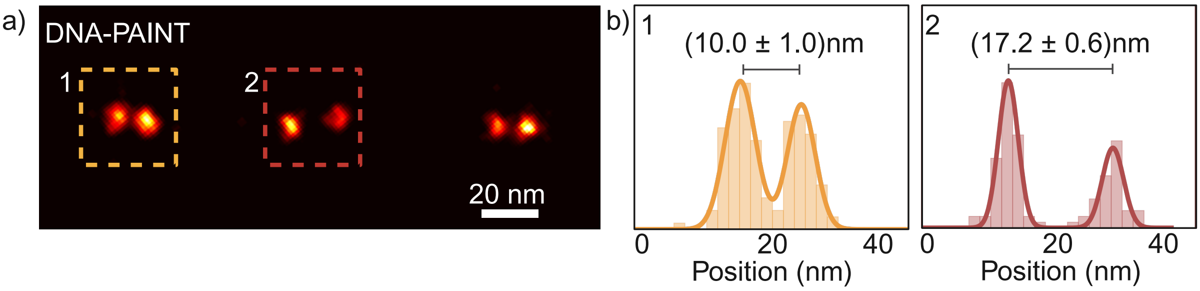


**Supplementary Figure 2:** (a) Super-resolved image of one DNA-origami structure measured using TIR microscopy showing the distances between docking sites for the center and corner pairs. (b) Histogram showing position distribution and measured distances of left (1) and central (2) docking sites pairs of (a). DNA origami sequence is shown in the methods, along with sample preparation, and imaging conditions are described in Supplementary Table 1. $\sigma_{SMLM}$: 1.0 nm. NeNa value: 2.7 nm. DNA origami sample imaging was repeated in two independent experiments.

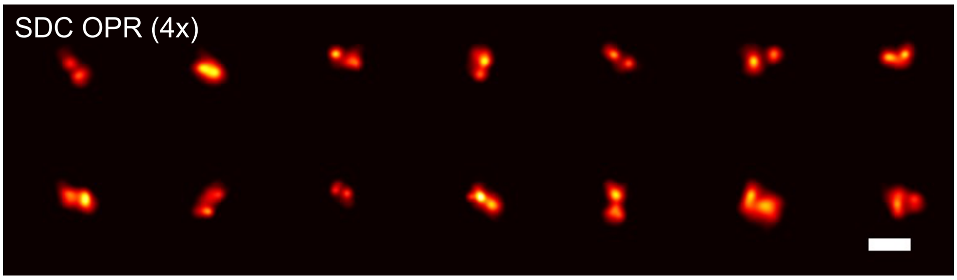


**Supplementary Figure 3:** Representative DNA-PAINT images of DNA-docking pairs spaced by 6 nm apart within DNA origami structures, acquired using 4× magnification with spinning disk confocal with optical photon reassignment (SDC-OPR) and deconvolution as described in Methods. DNA origami sequence is shown in Supplementary Table 2 and imaging conditions are described in Supplementary Table 1. Scale bar is 20 nm. $\sigma_{SMLM}$: 2.8 nm. NeNa value: 3.1 nm. DNA origami sample imaging was repeated in two independent experiments.

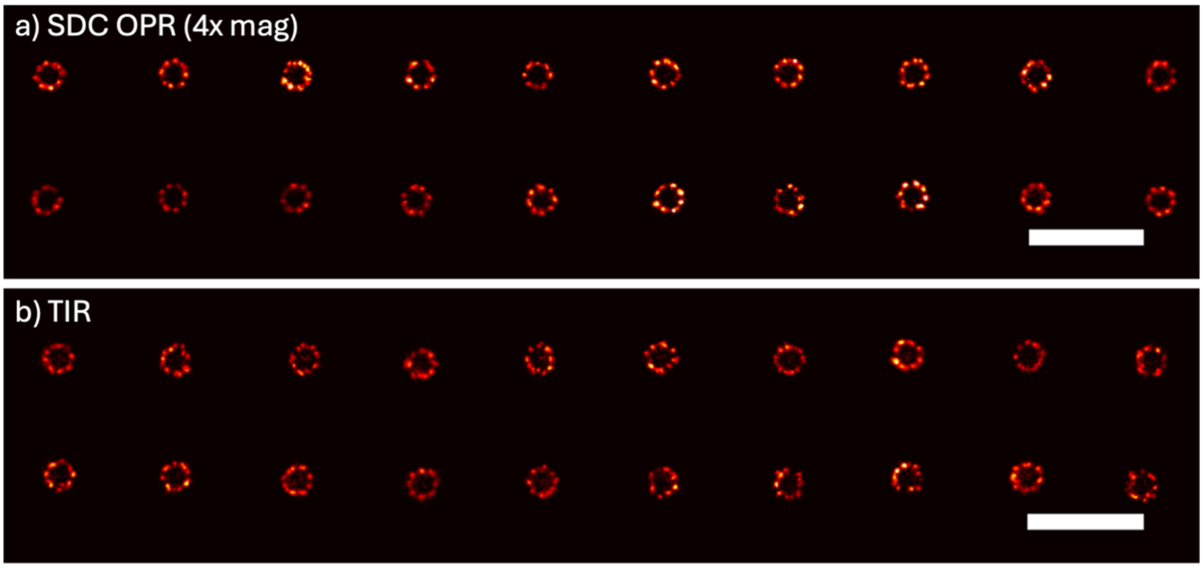


**Supplementary Figure 4:** NPC in U2OS cells imaged via Nup96, tagged with monomeric enhanced green fluorescent protein (mEGFP) and labelled with DNA-conjugated anti-GFP nanobodies for different excitation illumination. Sample preparation is described in the methods and imaging conditions are summarized in Supplementary Table 1. (a) Imaged with 4× magnification using spinning disk confocal with optical photons reassignment (SDC-OPR), with deconvolution as described in methods, showing 20 representative NPCs from the field of view. $\sigma_{SMLM}$: 3.3 nm. NeNa value: 4.4 nm. (b) Imaged with total internal reflection (TIR) illumination showing 20 representative NPCs from the field of view. $\sigma_{SMLM}$: 2.7 nm. NeNa value: 4.9 nm. Scale bar is 500 nm. Nup96 imaging was repeated in three independent experiments.

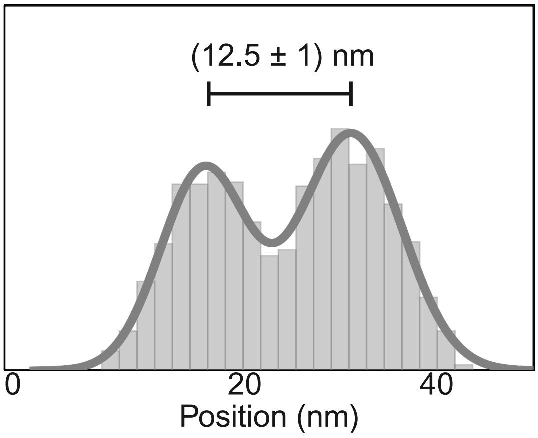


**Supplementary Figure 5:** Cross-sectional histogram of average protein pairs (*n* = 12 pairs) of Nup96 proteins in single symmetric centers measured using TIR microscopy, showing a 12.5 nm distance between single proteins.

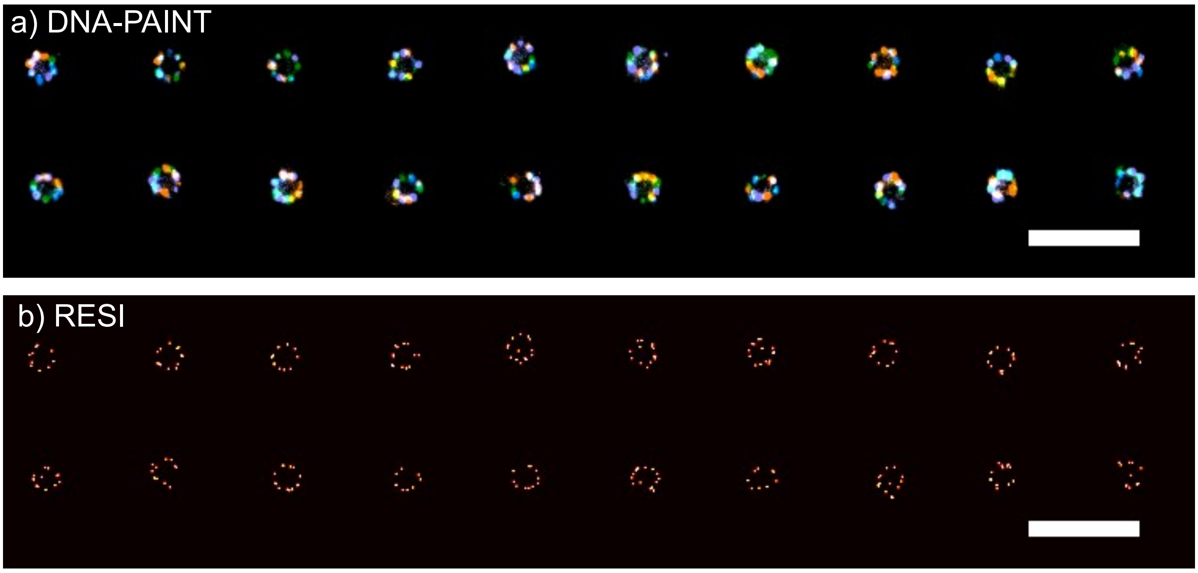


**Supplementary Figure 6:** RESI imaging of NPC in U2OS cells imaged via Nup96, tagged with monomeric enhanced green fluorescent protein (mEGFP) and labelled with four DNA-conjugated anti-GFP nanobodies for RESI imaging. Sample preparation is described in the methods and imaging conditions are summarized in Supplementary Table 1. (a) 4-color DNA-PAINT data prior to RESI analysis. Colors represent sequential imaging rounds. (b) Final resolved resolution enhancement by sequential imaging (RESI) data. Scale bar is 500 nm. $\sigma_{SMLM}$: 2.9 nm. NeNa value: 4.6 nm. Nup96 imaging was repeated in three independent experiments.

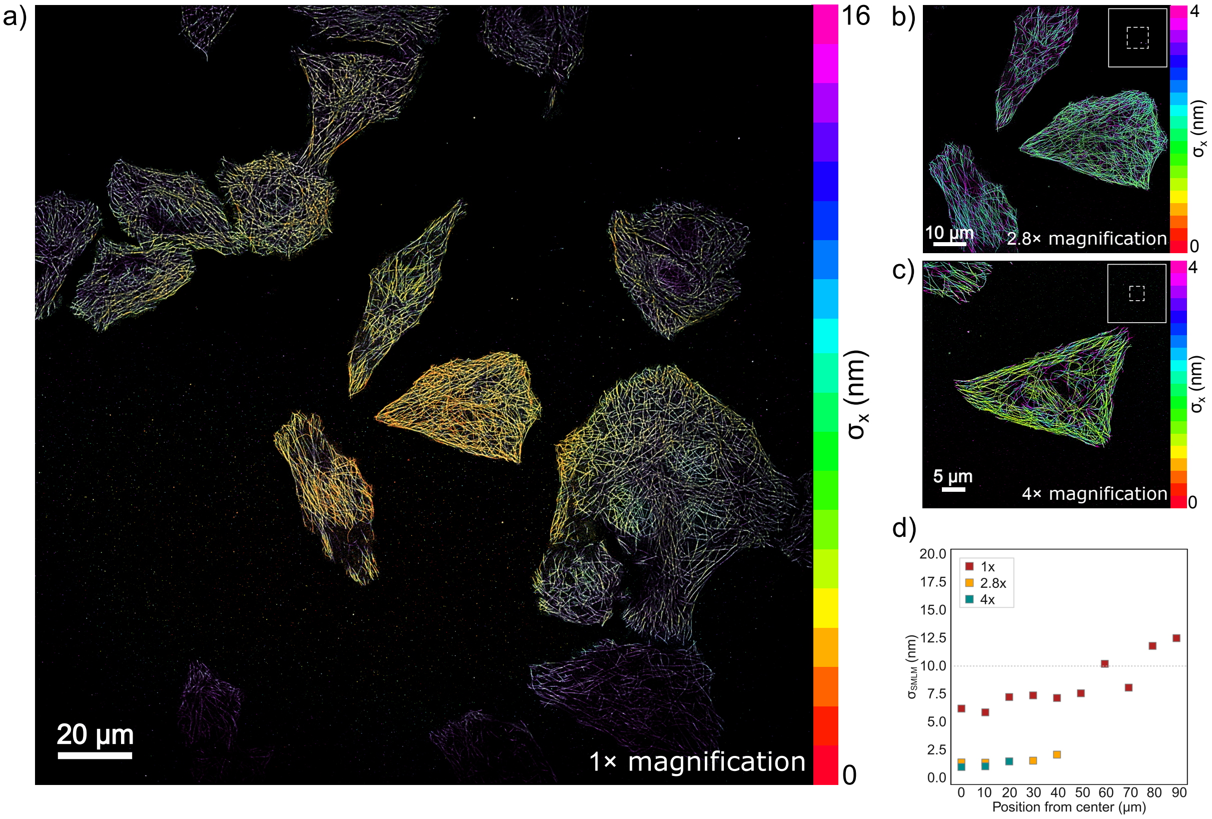


**Supplementary Figure 7:** DNA-PAINT overview of microtubules in HeLa cells across the FOV for different microscope magnifications where color represents the localization precision to evaluate the homogeneity across the whole FOV. Square inset in each image shows the area of illumination compared with the large FOV of 1× magnification. (a) 1× magnification, FOV 211 µm × 211 µm$^2$. $\sigma_{SMLM}$: 8.0 nm. NeNa value: 22.6 nm. (b) 2.8× magnification, FOV 76 µm × 76 µm$^2$. $\sigma_{SMLM}$: 1.6 nm. NeNa value: 4.2 nm. (c) 4× magnification, FOV 53 m × 53 µm$^2$. $\sigma_{SMLM}$: 1.7 nm. NeNa value: 3.7 nm. (d) Localization precision ($\sigma_{SMLM}$) measured at different distances (in µm) from the center of the FOV for 1× (red), 2.8× (yellow) and 4× (green) magnifications. For (a-c) color represents z position. Microtubule imaging was repeated in three independent experiments.

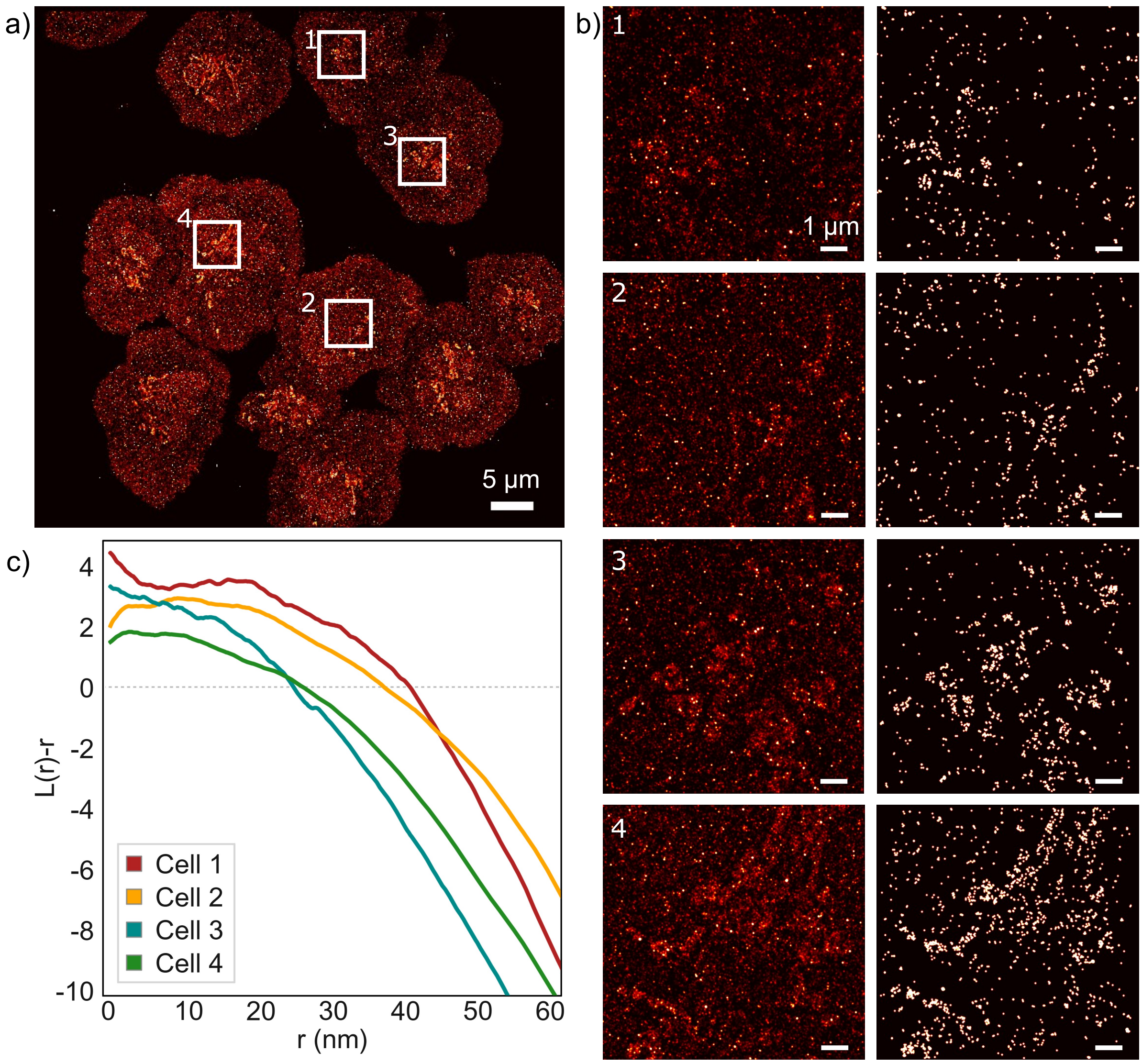


**Supplementary Figure 8:** DNA-PAINT imaging and cluster analysis of T cell receptor (TCR) proteins in Jurkat T cells activated for 10 min on glass coated with anti-CD3 and anti-CD28 antibodies. (a) DNA-PAINT image of TCR in Jurkat T cells acquired using SDC-OPR imaging at 2.8× magnification, capturing over ten cells within a single acquisition. Field of view (FOV): 76 × 76 µm². Scale bar = 5 µm. Single-molecule localization precision ($\sigma_{SMLM}$): 10 nm. NeNa value: 11 nm. (b) Zoom-in regions from (a) (top row), showing DNA-PAINT single-molecule localization maps corresponding to the white boxed areas. The bottom row displays reconstructed protein density maps generated using the qPAINT analysis pipeline (see Methods). Scale bar = 1 µm. (c) TCR clustering toward the central region of the T cell surface is crucial for activation and the initiation of downstream responses, including proliferation and cytokine secretion. Ripley's K-function analysis, L(r)-r, of the selected regions quantifies TCR clustering, revealing heterogeneity in cluster density across different Jurkat T cells within the same FOV. Each color represents the results for different cells. These results demonstrate the capability of DNA-PAINT combined with SDC-OPR imaging to capture and quantify cellular heterogeneity by integrating large FOV imaging with high-resolution precision, providing key insights into the variability of T cell activation mechanisms. T cell imaging was repeated in three independent experiments.

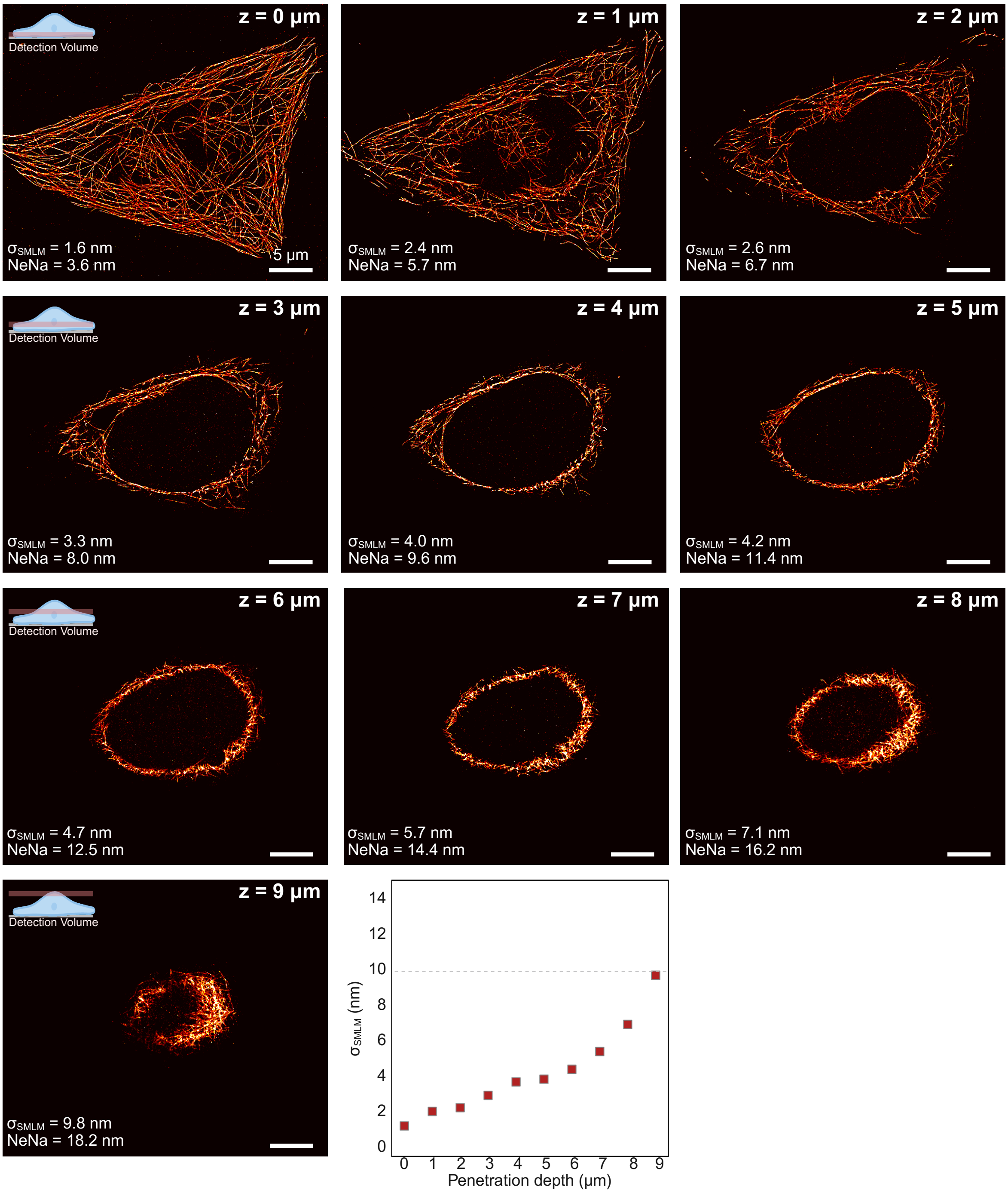


**Supplementary Figure 9:** DNA-PAINT z-stack sequence images of microtubules in HeLa cells, labelled with a-tubulin antibody, from z = 0 µm up to 9 µm with z-step size of 1 µm. Imaged with 4x magnification using SDC-OPR and deconvoluted as described in the methods. Localization precision ($\sigma_{SMLM}$) and nearest-neighbor-based metric (NeNa) values are shown for each respective plane and depicted. Sample preparation is described in the methods and imaging conditions are summarized in Supplementary Table 1. Scale bar is 5 µm. Microtubule imaging was repeated in three independent experiments. This figure was created with BioRender.com

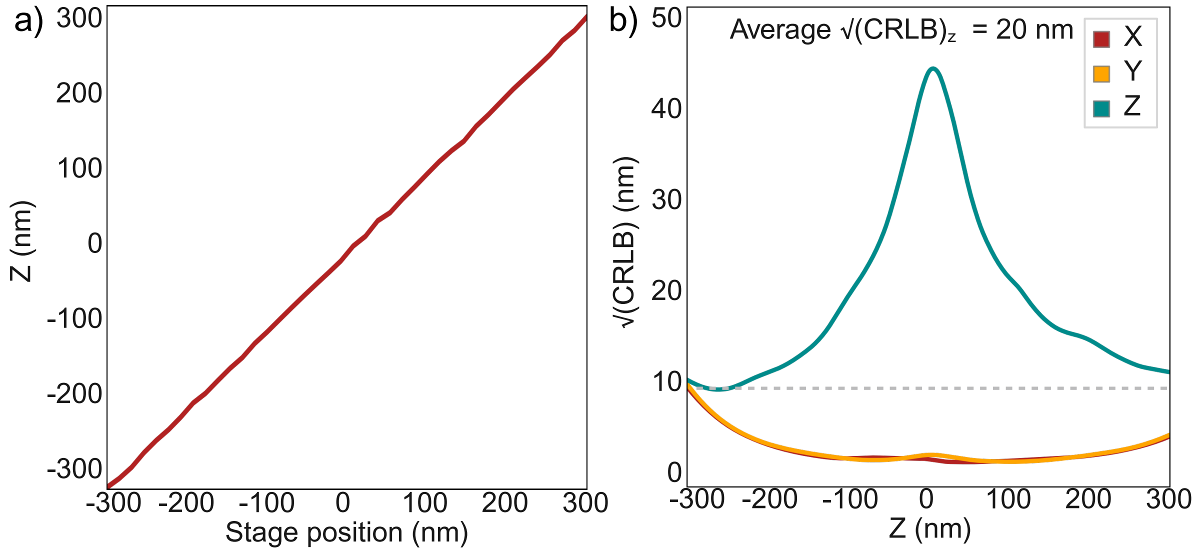


**Supplementary Figure 10:** *z* calibration data for 3D super-resolution experiments. (a) cSpline 2D interpolated PSF model was derived by imaging fluorescent beads (n = 8) with respect to stage position following the method described in Ref[25]. (b) Localization precision in 3D estimated by the Cramér–Rao lower bound (CRLB) at different axial positions for *x* (red) , *y* (yellow) and *z* (green)..

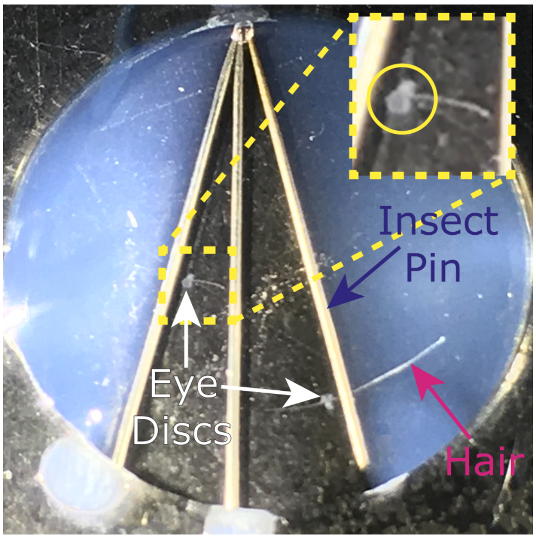


**Supplementary Figure 11:** Modified 35 mm glass bottomed dish used for imaging. Insect pins, hairs and 3rd instar Drosophila eye discs are indicated with blue, pink and white arrows respectively. A magnified eye disc is shown in the inset (yellow dashed square), circled in yellow.

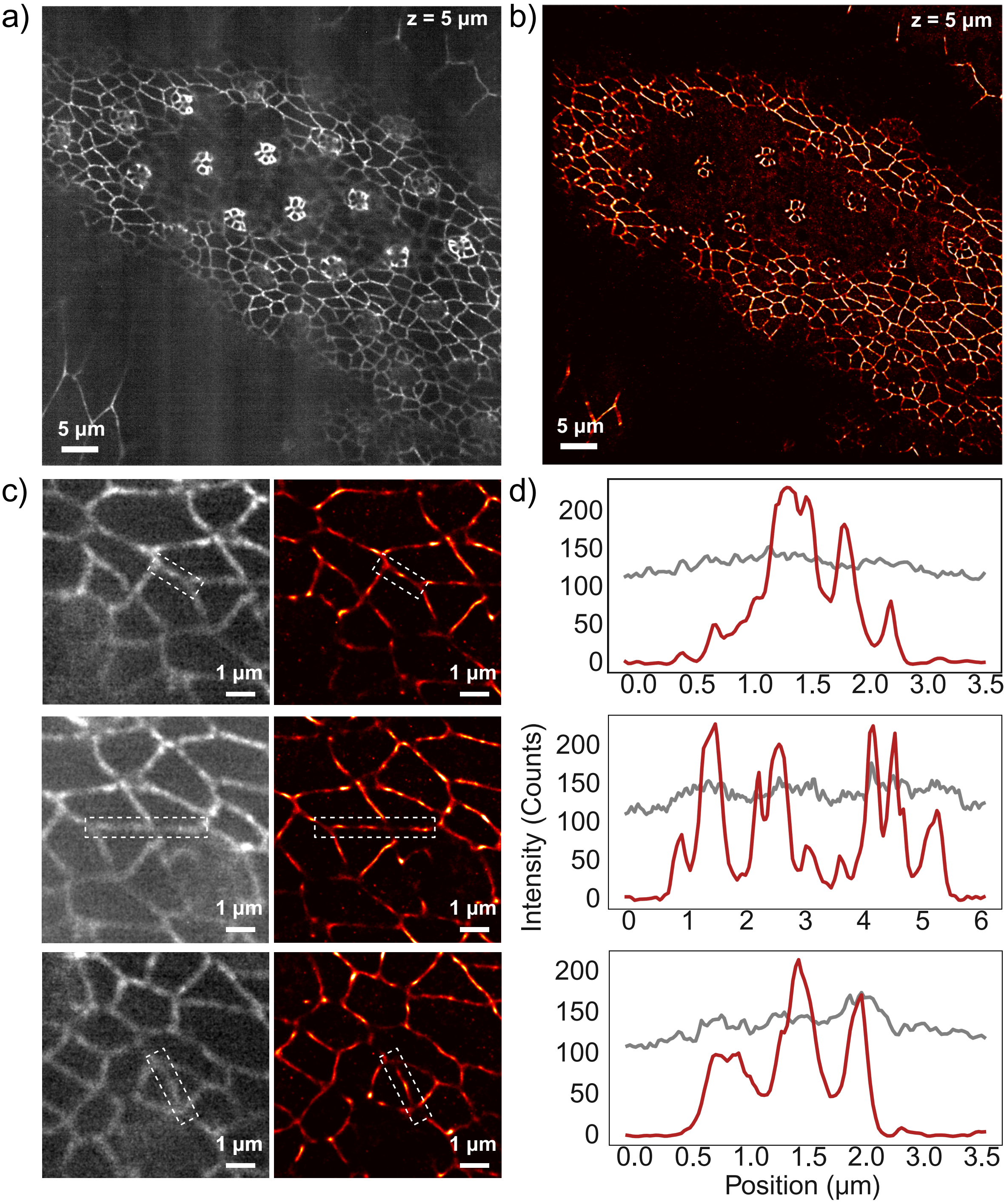


**Supplementary Figure 12:** Full-field view of (a) GFP-tagged and (b) DNA-PAINT imaging of E-cadherin within the *Drosophila* eye imaginal disc at a 5 µm penetration depth. $\sigma_{SMLM}$: 11.6 nm. NeNa value: 23.3 nm. (c) Zoomed-in 3.5 × 3.5 $\mu m^2$ regions for GFP (left) and DNA-PAINT (right). (d) Intensity plot profiles of the highlighted regions in (c) for GFP (gray) and DNA-PAINT (red), demonstrating the enhanced resolution achieved with DNA-PAINT compared to the diffraction-limited GFP channel. This improved resolution enables more accurate quantification of E-cadherin foci dimensions.

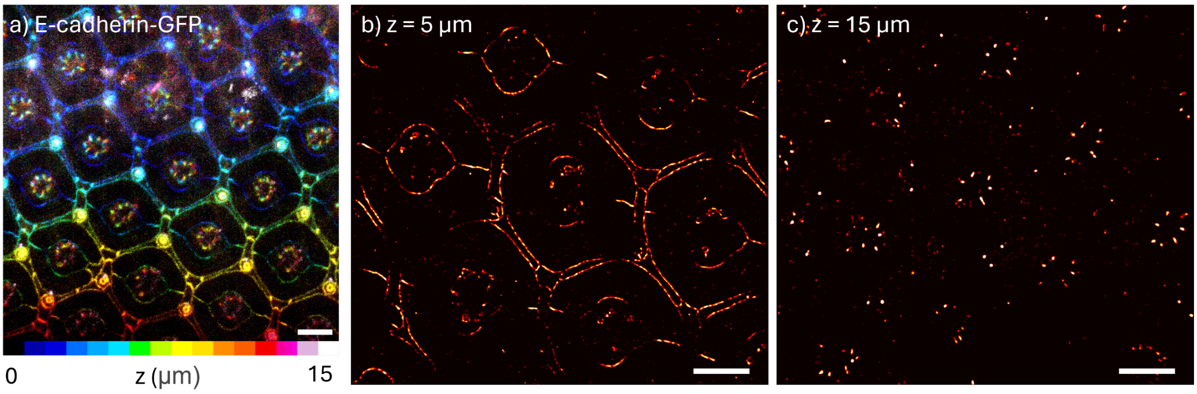


**Supplementary Figure 13:** (a) SPC-OPR 3D projection image of the GFP staining of E-cadherin for a total penetration depth of 15 μm, with 0.5 μm z-step, with color representing the *z* position. (b,c) SDC-OPR DNA-PAINT images of E-cadherin within the *Drosophila* pupal fly retina at 5 μm ($\sigma_{SMLM}$: 10.6 nm. NeNa value: 9.8 nm) and 15 μm ($\sigma_{SMLM}$: 26.4 nm. NeNa value: 21.3 nm), respectively. E-cadherin tagged with GFP was imaged using DNA-labelled anti-GFP antibodies, as described in the Methods section. Scale bar: 7 μm. All *Drosophila* imaging was repeated in three independent experiments

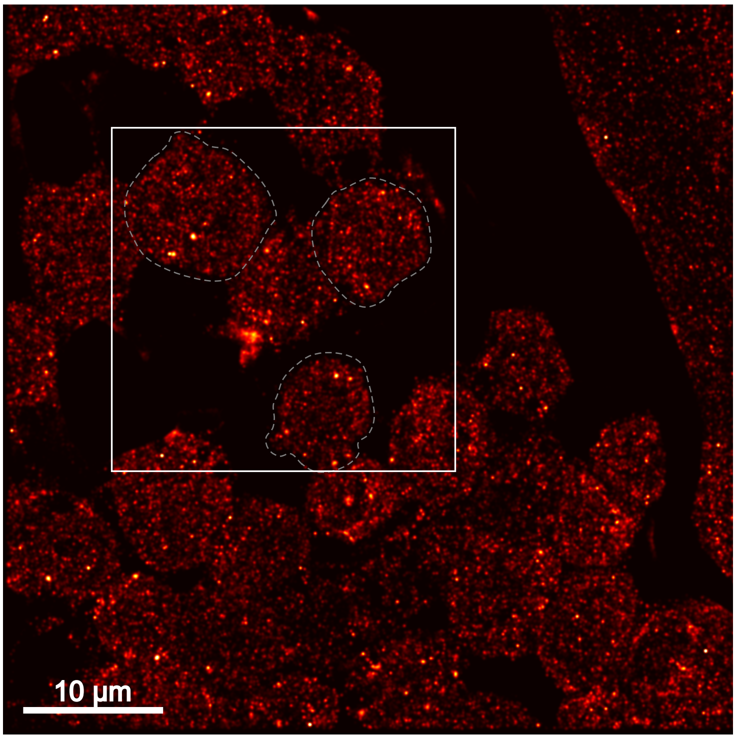


**Supplementary Figure 14:** Full Field of view of Collagen-IV imaging of the basement membrane. White rectangle points the area shown in Fig. 2i and 2 examples cells are outlined. Imaged with 4x magnification using SDC-OPR and deconvoluted as described in the methods. Sample preparation is described in the methods and imaging conditions are summarized in Supplementary Table 1. $\sigma_{SMLM}$: 4.6 nm. NeNa value: 19.6 nm. All *Drosophila* imaging was repeated in three independent experiments.

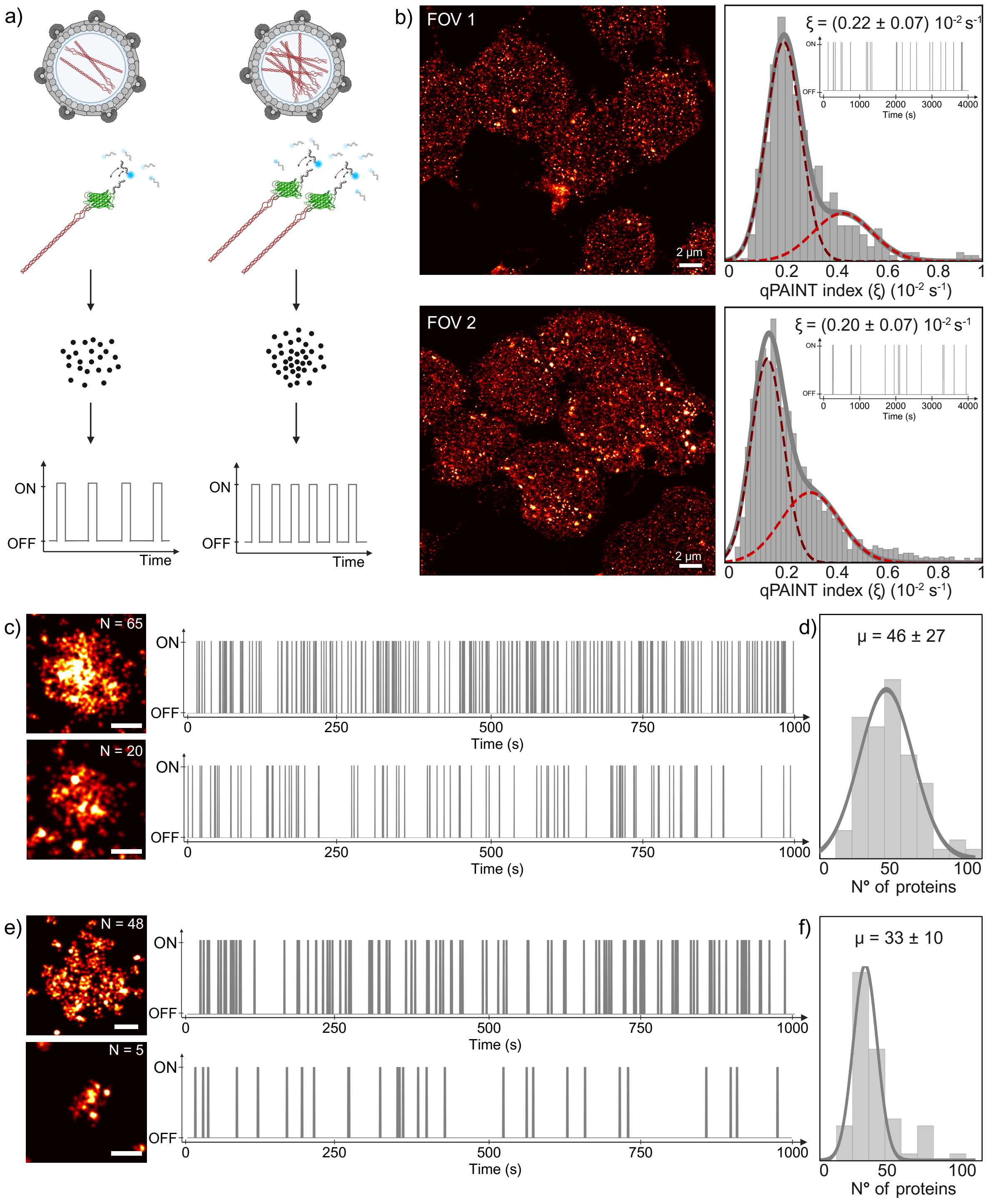


**Supplementary Figure 15:** Quantitative-Paint (qPAINT) calibration. (a) In DNA-PAINT, fluorescently tagged imager strands transiently bind to complementary docking strands anchored to a target—in this case, GFP-Collagen-IV proteins within vesicles. The fluorescence intensity traces generated from each vesicle's localization clusters over time exhibit distinct on- and off-time patterns, enabling the estimation of the number of molecules within a non-resolvable protein cluster —in this case a collagen vesicle. By determining the binding frequency of a single protein, the number of proteins in different clusters can be estimated by calculating the ratio of their binding frequencies. (b) Left: DNA-PAINT Collagen-IV imaging of the basement membrane for two independent field-of-views (FOVs). Right: Histogram of qPAINT indexes ($\xi$) for GFP:Collagen-IV proteins for each FOV. The fit to a sum of two Gaussian functions is shown as solid lines, whereas each component is shown as a dashed line. This fitted value $\xi$ was then used to calculate the number of proteins per cluster in the image. Inset shows blinking kinetics of an example single protein cluster. (c) Zoomed-in views of two example vesicles from FOV1, containing 65 and 20 proteins, respectively, along with their corresponding time traces, where the different binding frequency can be seen. (d) Histogram of proteins per vesicles for FOV1 showing the distribution average value ($\mu$). (e) Zoomed-in views of two example vesicles from FOV2, containing 48 and 5 proteins, respectively, along with their corresponding time traces, where the different binding frequency can be seen. (f) Histogram of proteins per vesicles for FOV2 and distribution average value ($\mu$). Panel (a) was created using BioRender.com

### Supplementary Tables

**Supplementary Table 1:** Summary of imaging conditions used for SDC, SDC-OPR and TIR DNA-PAINT with their respective datasets. Showing exposure time (ms), number of frames, power (both mW and W/cm$^2$), imager solution and buffer used.

| Data | Setup | Exposure time (ms) | Number of frames | Power (at objective) | Imagers | Buffer |
|---|---|---|---|---|---|---|
| Fig. 1c, SI Fig 1a | SDC | 300 | 15000 | 1.75 mW (4x mag) | R1 and R4 (1 nM both) | B+ (with PCA, PCD, Trolox) |
| Fig. 1d, Fig. 1f~~e~~ (top), SI Fig. 1b | SDC-OPR | 300 | 15000 | 1.75 mW (4x mag) | R1 and R4 (1 nM both) | B+ (with PCA, PCD, Trolox) |
| Fig. 1e, SI Fig. 3 | SDC-OPR | 300 | 17000 | 1.75 mW (4x mag) | R1 and R4 (1 nM both) | B+ (with PCA, PCD, Trolox) |
| Fig. 1f~~e~~ (center), SI Fig. 1c | SDC-OPR | 300 | 15000 | 3 mW (2.8x mag) | R1 and R4 (1 nM both) | B+ (with PCA, PCD, Trolox) |
| Fig. 1fe (low), SI Fig. 1d | SDC-OPR | 300 | 15000 | 5 mW (1x mag) | R1 and R4 (1 nM both) | B+ (with PCA, PCD, Trolox) |
| Fig. 2a, Fig.2c, SI Fig. 4~~3~~a | SDC-OPR | 300 | 15000 | 1.75 mW (4x mag) | F3 (1 nM) | C+ (with PCA, PCD, Trolox) |
| Fig. 3a | SDC-OPR | 300 | 15000 | 1.75 mW (4x mag) | F2 (0.3 nM) R2 (0.1 nM) F3 (1nM) | C+ (with PCA, PCD, Trolox) |
| Fig. 3b, SI Fig. 6 | SDC-OPR | 300 | 17000 | 1.75 mW (4x mag) | R1, R2, R3, R4 (8 nM all) | C+ (with PCA, PCD, Trolox) |
| Fig. 4~~3~~a, SI Fig 7~~5~~a | SDC-OPR | 300 | 17000 | 5 mW (1x mag) | R2 (1nM) | C+ (with PCA, PCD, Trolox) |
| Fig. 4~~3~~b, Fig. 4~~3~~d, SI Fig. 7~~5~~c, SI Fig 9~~ 6~~ | SDC-OPR | 300 | 17000 | 1.75 mW (4x mag) | R2 (1nM) | C+ |
| Figure 5 (3D whole cell) | SDC-OPR | 300 | 20000 | 1.75 mW (4x mag) | F2 (0.5 nM) | C+ |
| Fig, 6~~4~~c, Fig. 6~~4~~d, SI Fig. 12 | SDC-OPR | 300 | 18000 | 1.75 mW (4x mag) | R1 (10 nM) | C+ (with PCA, PCD, Trolox) |
| Fig. 6~~4~~e, SI Fig. 14~~8~~ | SDC-OPR | 300 | 20000 | 1.75 mW (4x mag) | R1 (5nM) | C+ (with PCA, PCD, Trolox) |
| SI Fig. 1e, SI Fig. 2 | TIR | 100 | 20000 | 5.7 mW (600 W/cm$^2$) | R1 and R4 (1 nM both) | B+ (with PCA, PCD, Trolox) |
| SI Fig. 4~~3~~b | TIR | 100 | 20000 | 3 mW (300 W/cm$^2$) | F3 (1 nM) | C+ (with PCA, PCD, Trolox) |
| SI Fig. 7~~5~~b | SDC-OPR | 300 | 17000 | 3 mW (2.8x mag) | R2 (1 nM) | C+ (with PCA, PCD, Trolox) |
| SI Fig. 8 | SDC-OPR | 200 | 10000 | 3 mW (2.8x mag) | R2 (2 nM) | C+ |
| SI Fig. 13 | SDC-OPR | 300 | 17000 | 1.75 mW (4x mag) | R1 (10 nM) | C+ (with PCA, PCD, Trolox) |

**Supplementary Table 2:** List of modified staples for the different DNA-PAINT sites for DNA-origami with edge pairs separated by 10 nm. Blue bases represent spacer sequence before docking strand sequence, red sequence represents docking strand sequence for corresponding DNA-PAINT imager to bind to.

| Staple Name | Sequence |
|---|---|
| **PL3 R1** | ATGCTGATGTTGGGTTTAGCTTAGATTAAGAC*TTTCCTCCTCCTCCTCCTCCT* |
| **PL1 R4** | TGAGCGCTTAAGCCCATGGCATGATTAAGACT*CAACACACACACACACACACA* |
| **PC2 R1** | CAGAACCGCATGTACCGCTAAACAACTTTCAA*TTTCCTCCTCCTCCTCCTCCT* |
| **PC8 R4** | AGCTCAACGAACGAGTGGCAAGGCAAAGAATT*CAACACACACACACACACACA* |
| **PR7 R1** | AAATTGTTATGGTCATCTCTTCGCTATTACGC*TTTCCTCCTCCTCCTCCTCCT* |
| **PR4 R4** | AAGAACTCCAATATTAGTCACACGACCAGTA*CAACACACACACACACACACA* |
| **Biotin 1** | ACAGGAAGATTGTCCCCCTTATTCACCCTCATTTGTTTC **Biotin** |
| **Biotin 2** | GTTGATAGATATAAGCATAAGTATAGC **Biotin** |
| **Biotin 3** | AGAGTACTCACGCTAACCTTTAATTGC **Biotin** |
| **Biotin 4** | CACTAAAACACTCACGAACTAACACTAAAGT **Biotin** |
| **Biotin 5** | TCACGACGTTGGGCGCTTTGGTAAAAC **Biotin** |
| **Biotin 6** | CAGAGATAGCGATAGTGAATAACATAA **Biotin** |
| **ATTO 647N** | CCCCTACCGACAAAAGGTAATAAGAGAATATAAAGCCCC **ATTO 647N** |
| **ATTO 532** | CCCCCGCTGAGAGCCAGCAGGCCTGCAACAGTGCCACCCC **ATTO 532** |

**Supplementary Table 3:** List of modified staples for the different DNA-PAINT sites for DNA-origami with edge pairs separated by 6 nm. Blue bases represent spacer sequence before docking strand sequence, red sequence represents docking strand sequence for corresponding DNA-PAINT imager to bind to.

| Staple Name | Sequence |
|---|---|
| **PL1 R3** | TGAGCGCTTAAGCCCATGGCATGATTAAGACTTC*CTCTCTCTCTCTCTCTCTC* |
| **PL6 R4** | AATTGAGTAATATCAGAAAATAAACAGCCATACA*ACACACACACACACACACA* |
| **PR2 R4** | TCGTAATCATCCGCTCATGAATCGGCCAACGC*CAACACACACACACACACACA* |
| **PR7 R3** | AAATTGTTATGGTCATCTCTTCGCTATTACGCTC*CTCTCTCTCTCTCTCTCTC* |
| **Biotin 1** | ACAGGAAGATTGTCCCCCTTATTCACCCTCATTTGTTTC **Biotin** |
| **Biotin 2** | GTTGATAGATATAAGCATAAGTATAGC **Biotin** |
| **Biotin 3** | AGAGTACTCACGCTAACCTTTAATTGC **Biotin** |
| **Biotin 4** | CACTAAAACACTCACGAACTAACACTAAAGT **Biotin** |
| **Biotin 5** | TCACGACGTTGGGCGCTTTGGTAAAAC **Biotin** |
| **Biotin 6** | CAGAGATAGCGATAGTGAATAACATAA **Biotin** |
